# Stochastic modelling of cell differentiation networks from partially-observed clonal tracking data

**DOI:** 10.1101/2022.07.08.499353

**Authors:** L. Del Core, D. Pellin, M. A. Grzegorczyk, E. C. Wit

## Abstract

**Motivation:** Clarifying how hematopoietic stem cells differentiate into mature cell types is important for understanding how they attain specific functions and offers the potential for therapeutic manipulation. Over the past decades, clonal tracking has proven to be capable of unveiling population dynamics and hierarchical relationships in vivo. For this reason, clonal tracking studies are required for safety and long-term efficacy assessment in gene therapy. However, many standard clonal tracking studies consider only a subset of cell-types and are subject to noise.

**Results:** In this work, we propose a stochastic framework that investigates the dynamics of cell differentiation from typical clonal tracking data subject to measurement noise, false-negative errors, and systematically unobserved cell types. Our framework is based on stochastic reaction networks combined with extended Kalman filtering and Rauch-Tung-Striebel smoothing. Our tool can provide statistical support to biologists in gene therapy clonal tracking studies to better understand clonal reconstitution dynamics.

**Availability:** The stochastic framework is implemented in the 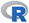 package Karen which is available for download at https://github.com/delcore-luca/Karen. The code that supports the findings of this study is openly available at https://github.com/delcore-luca/CellDifferentiationNetworks.

## 1 Introduction

Hematopoiesis is the process responsible for maintaining the number of circulating blood cells that are undergoing continuous turnover. This process has a tree-like structure with the root node constituted by Hematopoietic Stem Cells (HSC) [1; 2]. Each cell division gives rise to progeny cells that can retain the properties of their parent cell (self-renewal) or differentiate, “moving down” the hematopoietic tree [3–7]. As the progeny move further away from HSCs, their pluripotent ability is increasingly restricted. Clarifying how HSCs differentiate is essential for understanding how they attain specific functions and offers the potential for therapeutic manipulation [8]. Several mathematical models have been proposed to describe hematopoiesis in-vivo. One of the first stochastic models of hematopoiesis was introduced in the early ‘60s [9] suggesting that it is the population as a whole that is regulated rather than individual cells that behave stochastically, and control mechanisms act by varying the cell division and death rates.

More recently, [10–16] analyze data generated by using the most advanced lineage tracing protocols using novel statistical models. Still, to the best of our knowledge, none of the already existing tools considers the presence of false-negative clonal tracking errors. In addition to completely missing cell types, clonal tracking data are characterized by scattered detection (recapture) of clones due to either threshold detection failure or false-negative errors [17]. Usually, threshold detection failure is addressed by assuming that all the missing clone observations correspond to minimal clones and, therefore, set to zero. This hypothesis is too restrictive because it does not take into account other technical sources of false-negative errors, such as low-informative sample replicates [18]. It has also been shown that false-negative errors strongly depend on calling pipeline parameters, as well as read coverage [19]. The false-negative diagnosis rate is poorly understood for many NGS applications and is challenging to measure without the use of well-characterized reference standards [20]. The standardization of sequencing coverage depth has also been used to minimize the probability of false-positive and false-negative results. However, there is no consensus on the minimum coverage depth that clonal tracking data have to comply with, creating heterogeneity in the quality of data generated by the different laboratories [21].

We propose a stochastic framework to investigate haematopoiesis while cautiously treating all the undetected values as latent states. More precisely, we describe cell differentiation using stochastic quasi-reaction networks (SqRNs), a framework that allows to (i) model a network of stochastically interacting nodes using an Ito-type SDE formulation, and (ii) describe the dynamics of transition between different states (cell types) in terms of a set of reactions whose rates are unknown. Then we combine SqRNs with extended Kalman filtering (EKF) and Rauch-Tung-Striebel (RTS) smoothing. In particular, we (i) provide an expectation-maximization algorithm to infer the unknown parameters; (ii) extensively test the method on several simulation studies (iii) applied our framework to four in-vivo high dimensional clonal tracking data sets, to compare different biologically plausible models of cell differentiation. A flowchart of the analysis performed in this work is shown in Figure 1.

**Fig. 1:**
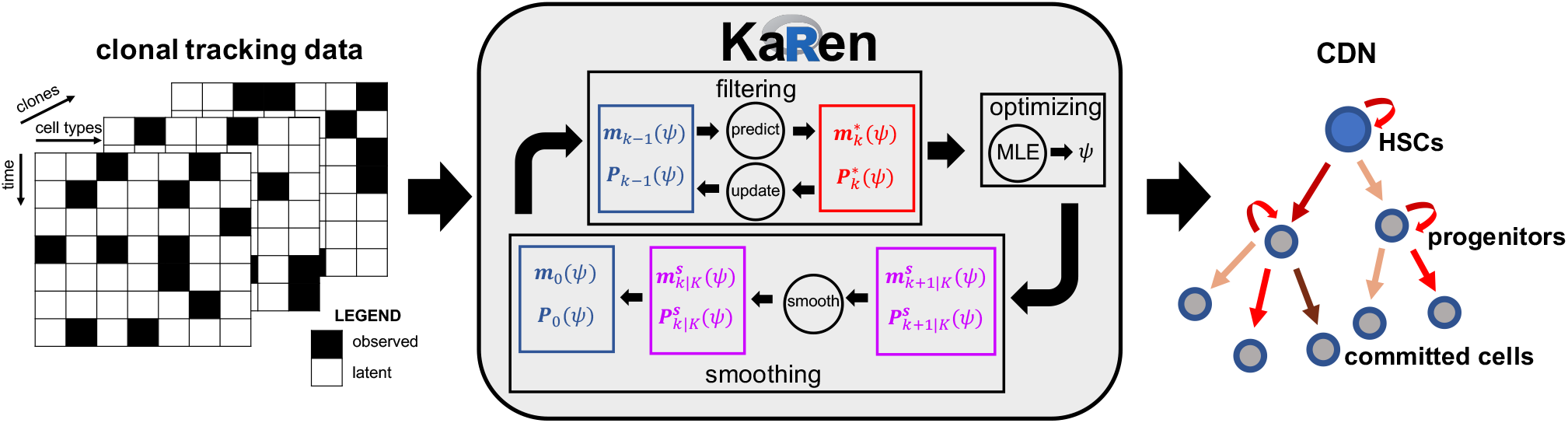
Schematic representation of the analysis flow: A three-dimensional clonal tracking dataset with partially-observed cells (left panel) is received as input from our proposed stochastic framework Karen (middle panel). It mainly consists in three parts, such as a filtering step, an optimization (maximum likelihood) step, and a smoothing step which are executed iteratively until a convergence is reached on the unknown vector parameter *ψ*. Finally, a cell differentiation network (CDN) is returned from Karen, where each arrow is directed and weighted according to the estimated parameters (right panel).

## 2 Methods

### 2.1 Karen: Kalman Reaction Networks

We consider a non-linear continuous-discrete state space model (CD-SSM) whose dynamic component is represented by the local linear approximation (LLA) of a stochastic quasi-reaction network [22] defined by a *n*×*J* net effect matrix *V*, a *p*×1 vector parameter *θ* and a *J* ×1 hazard vector *h*(*x*; *θ*) for a *n*-dimensional counting process {*x*(*t*)|*x*(*t*) ∈ ℕ^*n*^}_*t*_ (see Section S.1 from Supplementary Information for details). For the measurement function *g*_*k*_(*x*(*t*_*k*_), *r*_*k*_) we use a time-dependent selection matrix *G*_*k*_ ∈ **01**^*d×n*^ (the set of all *d* × *n* binary matrices) which selects only the measurable particles of *x*(*t*_*k*_) with an additive noise *r*_*k*_ whose covariance matrix is time-dependent and defined as

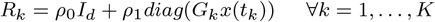

where *ρ*_0_ and *ρ*_1_ are free parameters which we infer from the data, and *diag*(·) is a diagonal matrix with diagonal equal to its argument. Therefore

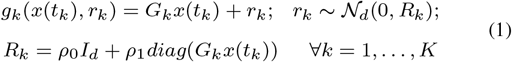

In the following *x*_*t*_ is a shorthand notation for *x*(*t*). Under these assumptions the CD-SSM of Eq. (1) from the **Supplementary Information** reduces to

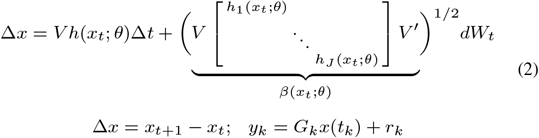

where

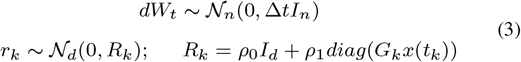

Assuming *x*(*t*_0_) ∼ 𝒩_*n*_(*x*(*t*_0_)|*m*_0_, *P*_0_) as prior distribution for *x*(*t*) at *t* = *t*_0_, the prediction step of Eq. (6) from the **Supplementary Information** reduces to

#### 1. Prediction step

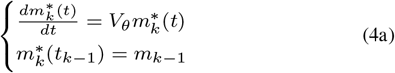

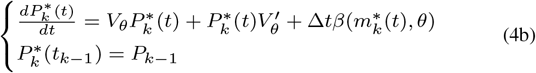

where we used the fact that for a set of reactions involving only one particle of *x* as reagent, which is the case for our cell differentiation networks, the mean drift *V h*(*x*_*t*_; *θ*) reduces to *V*_*θ*_*x*_*t*_, where the definition of *V*_*θ*_ depends on *V* and *h*(*x*^*t*^; *θ*). The solutions of (4) are given by

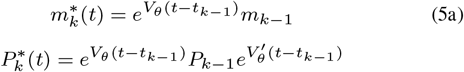

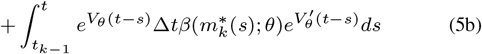

The solution for 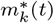 is obtained by applying the integrating factor method [23] to the initial value problem (4a) using an integrating factor 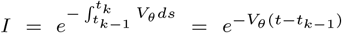. The solution for 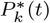 is obtained by applying the well-known solution formula for a differential Sylvester equation [24] to the system (4b). The corresponding update step is defined as follows

#### 2. Update step

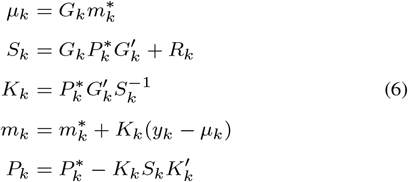

where 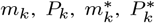, *μ*_*k*_ and *S*_*k*_ depend on both *θ, ρ*_0_ and *ρ*_1_. Finally, following Eq. (8) - (10) of the **Supplementary Information**, the optimization and smoothing steps are defined as

#### 3. Optimization step

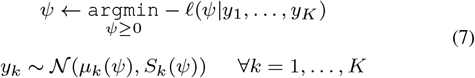

#### 4. Smoothing step

We use the Rauch-Tung-Striebel Smoothing algorithm (RTS) [25] and we estimate the first two-order moments of *p*(*x*_*k*_|*y*_1:*K*_, *θ, ρ*_0_, *ρ*_1_) as

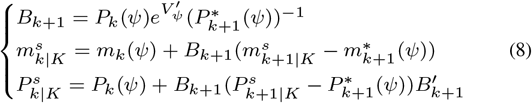

where *e*^(*·*)^ is the matrix exponential operator, *ψ* = (*θ, ρ*_0_, *ρ*_1_), and the values of 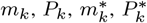, are the ones obtained from the filtering (prediction and update) steps. In order to run the optimization step using a gradient-based method (e.g. Newton-Raphson) we need to compute the gradient 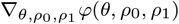 of the energy function *φ*(*θ, ρ*_0_, *ρ*_1_) which is defined by

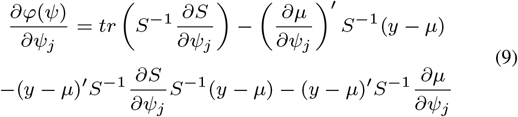

where

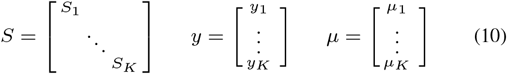

This requires, at every time point *k, p* + 2 more prediction and update steps in order to compute the 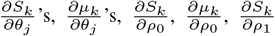 and 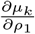, where *p* is the dimension of *θ*. These are obtained by deriving the equations in (4) and (6) w.r.t. *θ, ρ*_0_ and *ρ*_1_, as shown in Section S.2.1 of **Supplementary Information**. All the results obtained from every prediction/update step at each time point *t*_*k*_, along with the corresponding derivatives, are then used to compute the energy function *φ*(*θ, ρ*_0_, *ρ*_1_) and its gradient which, in turn, are used for the optimization step. The proposed extended Kalman filter is summarised in Algorithm 1 from Section S.4 of **Supplementary Information**. The whole procedure returns the estimated parameters 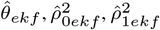 and the first two-order moments 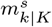 and 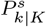 of the smoothing distribution *p*(*x*_*k*_|*y*_1:*K*_, *θ, ρ*_0_, *ρ*_1_) at every time point *t*_*k*_, *k* = 1, …, *K*. All the integrals involved for the computation of 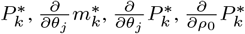 and 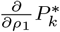 are estimated using a 3rd-order Gauss-Legendre method [26].

### 2.2 Stochastic formulation of clonal dynamics

We assume that the time counting process

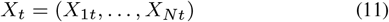

of *N* distinct cell types for a single clone evolves in a time interval (*t, t* + Δ*t*) according to a set of reactions and hazard functions defined as

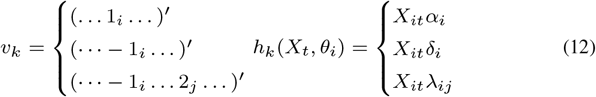

The hazard functions contain linear terms for duplication and death of cell *i* with positive rates *α*_*i*_ and *δ*_*i*_, and a linear term to describe cell differentiation from lineage *i* to lineage *j* with positive rate *λ*_*ij*_ for each i≠ *j* = 1, …, *N*. Finally, we use the compact matrix formulations

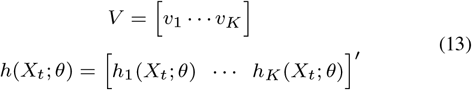

where *θ* is the vector of all the unknown parameters describing the dynamics. Since for our applications both the HSCs and the progenitors *P*_*i*_s are missing states, to help parameter inference of the state space model (2)-(3) combined with net-effect matrix and hazard vector (12) we assume the following conservation laws

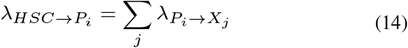

where *X*_*j*_ s are all the offspring cell types generated by *P*_*i*_.

### 2.3 Transition probabilities

Once the vector parameter *θ* is estimated for a particular model ℳ, we define the transition probability *p*_*ij*_ from cell type *i* to cell type *j* as

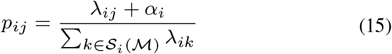

where 𝒮_*i*_(ℳ) is the set of all the possible target cell types associated to cell type *i* in the model ℳ.

### 2.4 Computational implementation

The stochastic framework is implemented in the 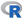 package Karen available at https://github.com/delcore-luca/Karen. Working examples showing the usage of the package are provided in Section S.7 of **Supplementary Information**.

## 3 Results

We use our stochastic framework to compare four different biologically-sustained models of hematopoiesis. The graphical representation of the candidate models is shown in Figure 2. For each candidate model, the stochastic differential equation formulation can be obtained from equations (12)-(13). Biological interpretation of the proposed models can be found in Section S.6 of the **Supplementary Information**.

**Fig. 2:**
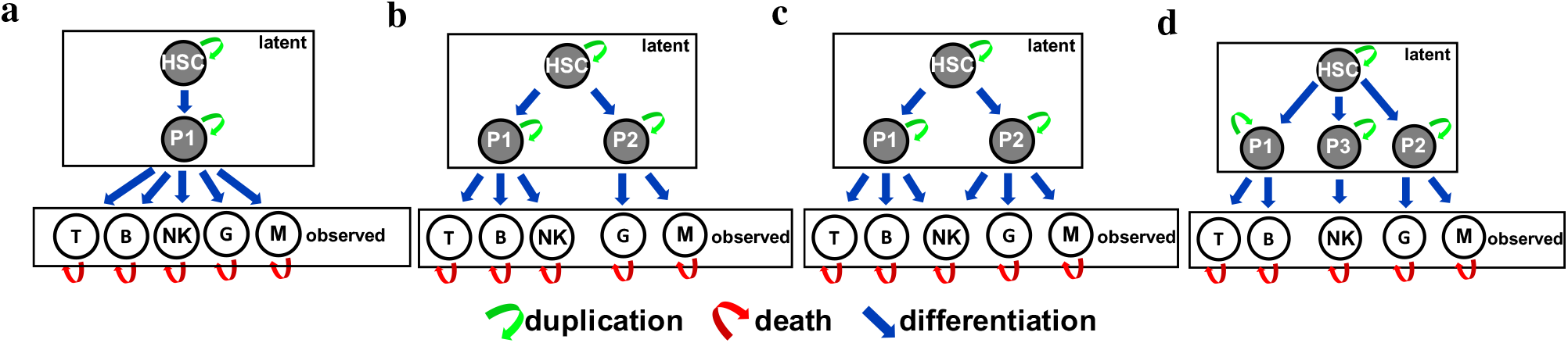
Graphical representation of the candidate models (a-d): Latent and observed cell types are indicated with grey and white nodes respectively. Red arrows denote a death move, green arrows indicate a duplication move, and blue arrows a differentiation move.

### Application to simulated data

We performed several simulations designed to test the proposed inferential procedure under different scenarios. The performance have been investigated by: (i) reducing the number of time points, (ii) reducing the fraction of clones recaptured across lineages and time, which is equivalent to increasing the rate of false-negative errors, (iii) increasing measurement noise, and (iv) selecting a cell differentiation structure among a set of candidates. Additional details on the simulation studies can be found in Section S.3 of the **Supplementary Information**. The results show the accurate recovery by the method of the true parameters, the first two-order smoothing moments, and the true generative model.

### Application to Rhesus Macaques study

We analyzed an in-vivo clonal tracking dataset previously used to investigate the hematopoietic reconstitution in Rhesus Macaques [27]. A pool of autologous CD34+ HSPCs barcoded by using lentiviral vectors have been transplanted in three myeloablated animals [28; 29]. Following engraftment, Granulocytes (G), Monocytes (M), T, B and NK cells were flow-sorted from peripheral blood (purity median 98.8%), and the majority of transduced cells contained only one barcode. Barcode retrieval by PCR was performed on purified hematopoietic lineage samples monthly for 4.5 months (ZG66), 6.5 months (ZH17), and 9.5 months (ZH33) [30]. Further details on transductions protocol and culture conditions can be found in the original paper study [27]. Although the sample DNA amount was maintained constant during the whole experiment (200 ng for ZH33 and ZG66 or 500 ng for ZH17), the sample collected resulted in different magnitudes of the total number of reads (see Table S.1 in **Supplementary Information**). This discrepancy makes samples not directly comparable across time and cell types. Therefore we rescaled the barcode counts as described in Section S.5 of the **Supplementary Information**. We report the rescaled cell counts, at the clonal level, in Figure 3. The total numbers of clones collected are 1165 (ZH33), 1280 (ZH17), and 1291(ZG66). We only focused on the top 1000 most recaptured clones across lineages and time to further remove bias.

**Fig. 3:**
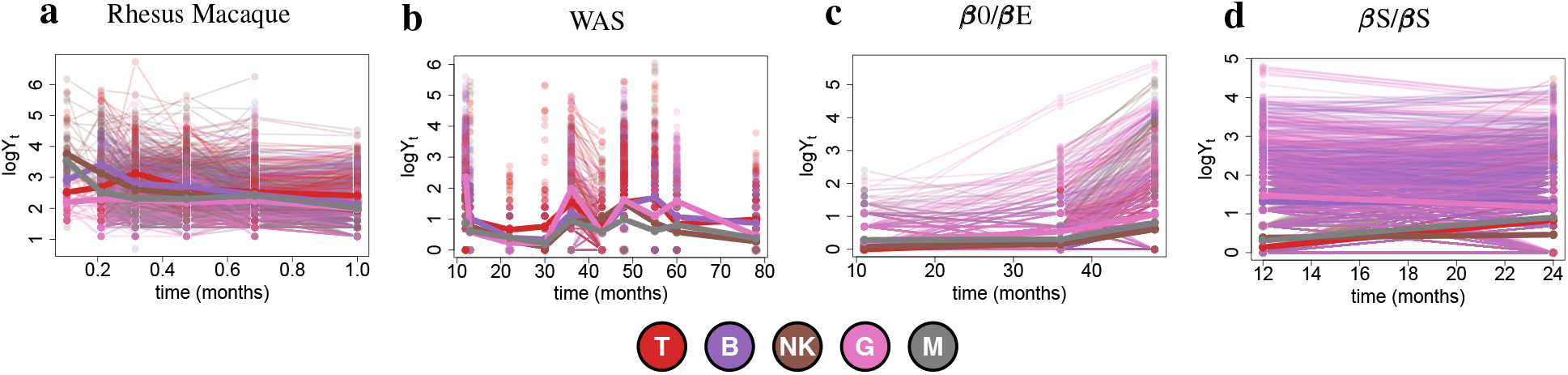
Clonal tracking data: Logarithmic clonal abundance (*y*-axis) over time (*x*-axis) in each cell lineage (colors) for the rhesus macaque study (a) and the clinical trials (b-d).

We fit the four candidate models on the clonal tracking data using Algorithm 1 from Section S.4 of **Supplementary Information**. We report the results in Figure 4 which shows, for each candidate model, the estimated cell differentiation network and the corresponding Akaike Information Criterion (AIC) [31] which we use as a measure of model selection. According to the AIC, model (c) is the one that best fits the clonal tracking data collected from the rhesus macaque study. This result suggests that the classical/dichotomic model (a) fails to describe adequately clonal dynamics in rhesus macaque, whereas the myeloid-based developmental model (c) better explains hematopoietic reconstitution. Therefore our proposed framework Karen clearly indicates that in primate hematopoiesis myeloid progenitors represent a prototype of hematopoietic cells capable to produce both myeloid G/M cells and NK cells.

**Fig. 4:**
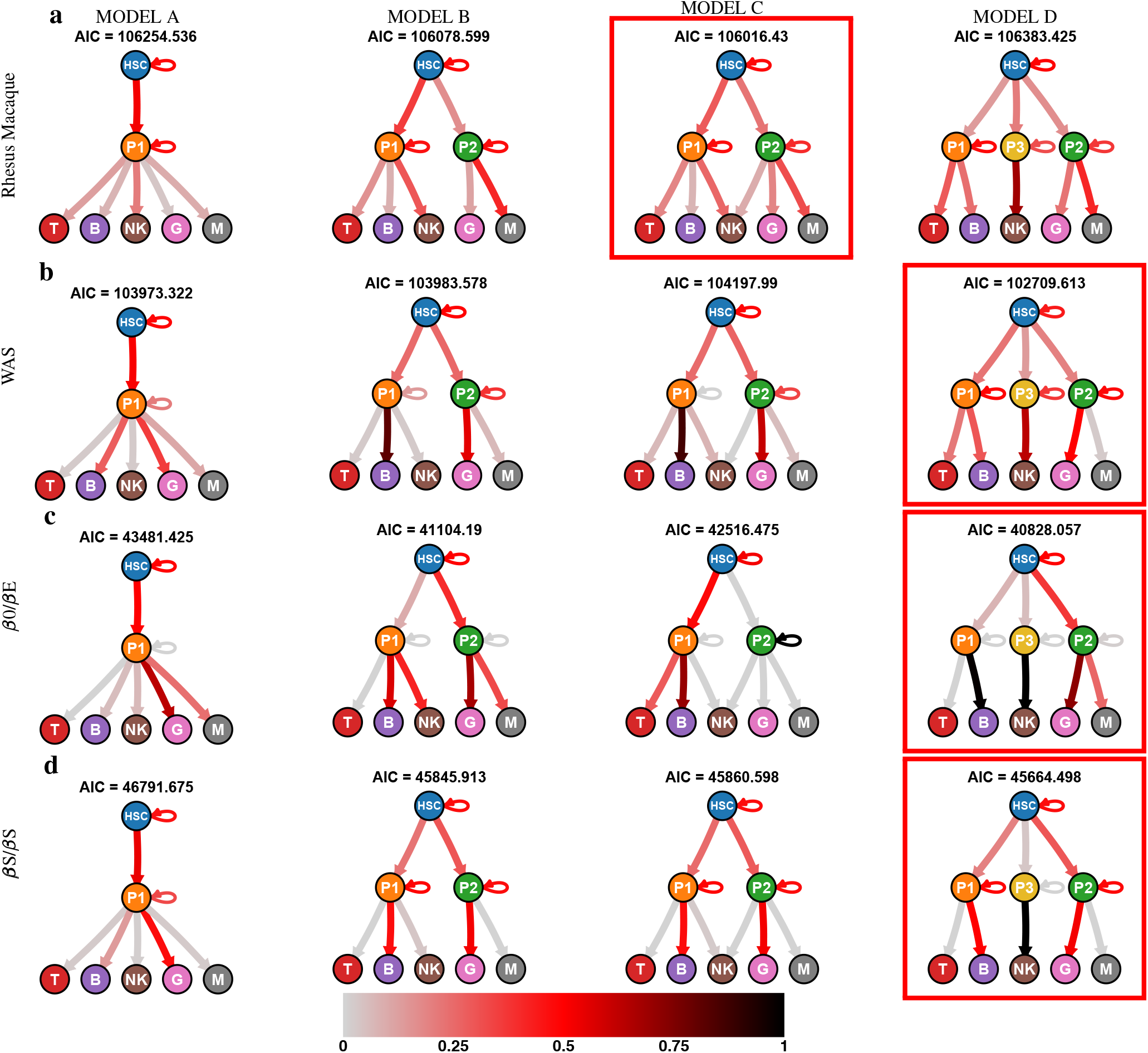
Results: Inferred cell differentiation networks for the candidate models (columns) for the rhesus macaque study (a) and the clinical trials (b-d). Each arrow is weighted and coloured according to the corresponding transition probability estimated with (15). For each model the AIC is reported and the best model is squared with a red box.

### Application to gene therapy clinical trials

Clonal tracking data derived from the analysis of samples isolated from six patients treated using HSPC-based gene therapy have been used to investigate human hematopoiesis dynamics. Five cell lineages (G, M, T, B, and NK) were collected longitudinally from the peripheral blood of four patients affected by Wiskott-Aldrich syndrome (WAS) [32], 2 patients with *β* hemoglobinopathy (1 with *β*S/*β*S sickle cell disease [33] and 1 with *β*0/*β*E *β* thalassemia [34]).

Details on procedures, gene therapy protocols, and normalization methods can be found in [32–35]. We report the clonal level logarithmic cell counts in Figure 3. Since data was already normalized to compensate for unbalanced sampling in VCN and DNA [32–35], we did not apply any further transformation. The total clones collected are 156654, 17273, and 230408, respectively, for WAS, *β*S/*β*S and *β*0/*β*E clinical trials. The following results derive from the analysis of the 1000 most recaptured clones in each clinical trial (top 250 clones per WAS patient).

The same four biologically motivated hematopoietic models (Figure 2) have been scored separately in each clinical trial using our stochastic framework Karen. We report the results in Figure 4 showing the estimated cell differentiation networks for each clinical trial. As a result, according to the AIC, model (d) is the one that always best fits clonal tracking data collected from each clinical trial, thus suggesting that a three-branches developmental model better explains hematopoietic reconstitution in humans after a gene therapy treatment. In particular, while lymphoid T/B and myeloid G/M develop in parallel trough separate branches from different progenitors, there is a third developmental branch for the lymphoid NK cells which is independent from the first two branches.

## 4 Discussion

We have proposed a novel stochastic framework for modeling cell differentiation networks from partially-observed high-dimensional clonal tracking data. Our model is able to deal with experimental clonal tracking data that suffers from measurement noise and low levels of clonal recapture due to either threshold detection failures or false-negative errors. Our framework extends stochastic quasi-reaction networks by introducing EKF and RTS components. We developed a tailor-made Expectation-Maximization (EM) algorithm to infer the corresponding parameters. Simulation studies have shown the method’s accuracy regarding inference of the true parameters, estimation of the first two-order smoothing moments of all the process states, and model selection. Simulation results indicated the method’s robustness in situations characterized by: the availability of a limited number of time points, limited clonal recapture, and high levels of measurement noise.

Although the gaussian assumption makes the analytical formulations of the likelihoods explicitly available, this approximation may become poor when the data contains outliers or shows non-gaussian behaviors. This limitation can be overcome by using a distribution-free approach, such as the Kernel Kalman Rule [36; 37]. Another limitation is that our framework considers reaction rates constant for the whole study period. Extensions that allow for modeling reaction rates as spline functions of time or depending on clinically relevant variables are within reach and will be the goal of future research.

The application of Karen on a rhesus macaque clonal tracking study unveiled for the lymphoid NK cells a different developmental pathway from the one detected for lymphoid T and B cells. That is, NK cells are produced by both myeloid and lymphoid progenitors P1 and P2. Results are consistent with the ones previously reported in [27] where the authors demonstrated the presence of distinct subpopulations within the NK lineage, potentially deriving from alternative maturation processes. Subsequently, we analyzed in-vivo clonal tracking data from three different clinical trials, showing consistency in the selected hematopoietic model structure across the clinical trials. Our stochastic framework can support biologists in understanding hematopoietic reconstitution and in designing tailor-made therapies to treat genetic disorders. Our model can be applied to different types of clonal tracking data, such as vector integration sites, clonal barcodes, and single cells methods. Applications in alternative contexts, such as the modeling of population dynamics, where similar issues about partial sampling and varying levels of measurement noise are present, could also be explored.

## Acknowledgements and Funding

This publication is based on work from COST Action CA15109 (COSTNET), supported by COST (European Cooperation in Science and Technology). E.C.W. acknowledges support from the Fondazione Leonardo (514.7.010.098-4) and funding from the Swiss National Science Foundation (SNSF 188534).

## Author Contributions

All authors contributed to analysing the data and writing the manuscript. L.D.C. designed and implemented the stochastic framework.

