## Supplementary Information for "Stochastic modelling of cell differentiation networks from partially-observed clonal tracking data"

### Supplementary Information for the paper "Stochastic modelling of cell differentiation networks from partially-observed clonal tracking data"

L. Del Core et al.


#### S.1 Stochastic quasi-reaction networks

##### S.1.1 Biochemical reactions

Stochastic quasi-reaction networks (S-QRN) allow to implement a particular class of stochastic differential equations that can be used to model biochemical reactions. More formally, let

$$Y_t = (Y_{1t}, \dots, Y_{Nt})$$

be a collection of molecules of $N$ different types observed at time $t$, and consider $K$ distinct (and competing) reactions

$$r_{j1}Y_1 + \cdots + r_{jN}Y_N \xrightarrow{\theta_j} p_{j1}Y_1 + \cdots + p_{jN}Y_N \qquad j = 1, \dots, K \tag{1}$$

each occurring with its own rate $\theta_j$. The coefficients $r_{ji}$'s defining the left-side of the reaction are called reagents and represent the minimum amount of molecules of type $i$ needed for the $j$-th reaction to occur. Similarly, the coefficients $p_{ji}$ defining the right-side of the reaction are called products and represents the amount of produced molecules of type $i$ after the $j$-th reaction is triggered. We assume that, if we observe $Y_0 = (r_{j1}, \dots, r_{jN})$ molecules at time $t = 0$, the $j$-th reaction will occur after

$$T_j \sim exp(\theta_j), \qquad j = 1, \dots, K$$

Namely, if exactly $r_{ij}$ molecules of each type $i$ would be present, then the $j$-th reaction can only take place in one way, with the exponential hazard rate $\theta_j$. The interpretation is that, after a waiting time $T_j$, $r_{ji}$ molecules of type $i$ collide with each other and produce $p_{ji}$ molecules of type $i$ ($\forall i = 1, \dots, N$), while the molecules move randomly in a hosting "cellular" environment. However, in general at time $t = 0$ we might observe $Y_{i0} \geq r_{ji}$ molecules of each type $i$ and, therefore, the $j$-th reaction can take place in a combinatorial number of ways leading to the following waiting time formulation

$$T_j \sim exp\underbrace{\left(\theta_j \prod_{i=1}^{N} \binom{Y_{i0}}{r_{ji}}\right)}_{h(Y_0,\theta)}, \qquad \texttt{where } \binom{x}{y} = 0 \texttt{ for } x < y \tag{2}$$

In this case, the effect will be that at time $t+T_j$ we have the following expression for the number of molecules of substrate $i$,

$$Y_{i,t+T_j} = Y_{it} + p_{ji} - r_{ji} = Y_{it} + v_{ji} \tag{3}$$

where $v_{ji} = p_{ji} - r_{ji}$ is the $j$-th net effect. More compactly, for a set of $K$ reactions and $N$ species, the molecular transfer from reagent to product species is a net change of

$$V = P - R$$

where $P = [p_{ji}]'$ denotes the $N \times r$ dimensional matrix of products, $R = [r_{ji}]'$ is the $N \times r$ dimensional matrix of reactants, and $V = [v_{ji}]'$ is an $N \times r$ dimensional matrix called net-effect matrix. Therefore, a S-QRN of $K$-distinct reactions is fully identified by a net-effect matrix $V$ and by the hazard vector

$$h(Y, \theta) = \begin{bmatrix} h_1(Y, \theta) & \cdots & h_K(Y, \theta) \end{bmatrix}'$$

##### S.1.2 Local linear approximation

We focus on the first two-order moments of the process, that is we consider the Ito equation

$$dY_t = \mu(dY_t; \theta)dt + \beta^{1/2}(dY_t; \theta)dW(t) \qquad dW(t) \sim N(0, dtI) \tag{4}$$

where $dY_t = Y_{t+dt} - Y_t$ is an infinitesimal time drift and $\mu(dY_t; \theta)$ and $\beta(dY_t; \theta)$ are called mean-drift and diffusion respectively. Given a $\sigma$-algebra $(\Omega, \mathcal{F}, \mathbb{P})$, the solution

$$Y : [0, +\infty) \times \Omega \to \mathbb{R}^N$$

of (4) is called Ito diffusion. Instead of finding the Ito diffusion itself, we focus on the first two-order moments $\mu(dY_t; \theta)$ and $\beta(dY_t; \theta)$ of the infinitesimal time drift $dY_t$ which can be approximated with the following Lemma and Proposition.

**Lemma 1.** *Given the hazard function as a limit of a conditional probability*

$$h(t) = \lim_{dt \to 0} \frac{\mathsf{P}(T < t + dt | T > t)}{dt}$$

*for small $dt$ the following approximation holds*

$$h(t)dt \approx \mathsf{P}(T < t + dt | T > t)$$

*Furthermore, the event $\{Y_{t+dt} - Y_t = V_{\cdot j}\} \equiv \{Y_{t+dt} = Y_t + V_{\cdot j}\}$ occurs with probability $\mathsf{P}(T_j < t + dt | T_j > t)$.*

**Proposition 1.** *An approximation of the mean drift $\mu(dY_t; \theta)$ and the diffusion $\beta(dY_t; \theta)$ for a small time increment $dt$ is given by*

$$\mu(dY_t; \theta) \underset{\texttt{small}\ dt}{\approx} Vh(Y_t, \theta) \tag{5}$$

$$\beta(dY_t; \theta) \underset{\texttt{small}\ dt}{\approx} V \underbrace{\begin{bmatrix} h_1(Y_t;\theta) & & \\ & \ddots & \\ & & h_K(Y_t;\theta) \end{bmatrix}}_{diag(h(Y_t, \theta))} V' \tag{6}$$

*Proof.*

$$\begin{aligned} \mu(dY_t; \theta) &\widehat{=} \lim_{dt \to 0} \frac{E[dY_t | Y_t, \theta]}{dt} \\ &= \lim_{dt \to 0} \frac{\sum_{j=1}^{K} \mathsf{P}(T_j < t + dt | T_j > t) V_{\cdot j}}{dt} \\ &\underset{\texttt{small}\ dt}{\approx} \lim_{dt \to 0} \frac{\sum_{j=1}^{K} h_j(Y_t, \theta) V_{\cdot j}}{dt} = Vh(Y_t, \theta) \end{aligned}$$

$$
\begin{aligned}
\beta(dY_t;\theta) &\widehat{=} \lim_{dt\to 0} \frac{Cov(dY_t|Y_t,\theta)}{dt} \\
&= \lim_{dt\to 0} \frac{E[dY_t dY_t'|Y_t;\theta] - E[dY_t|Y_t;\theta]E[dY_t|Y_t;\theta]'}{dt} \\
&\underset{\texttt{small } dt}{\approx} \frac{\sum_{j=1}^{K} V_{\cdot j}V_{\cdot j}' h_j(Y_t;\theta)dt - Vh(Y_t;\theta)h'(Y_t;\theta)V'dt^2}{dt} \\
&= \sum_{j=1}^{K} V_{\cdot j}V_{\cdot j}' h_j(Y_t;\theta) = V \begin{bmatrix} h_1(Y_t;\theta) & & \\ & \ddots & \\ & & h_K(Y_t;\theta) \end{bmatrix} V'
\end{aligned}
$$

□

Using previous results we can rewrite the Ito equation 4 in the following state-space formulation

$$
\begin{aligned}
dY_t = Vh(Y_t,\theta)dt + \left( V \begin{bmatrix} h_1(Y_t;\theta) & & \\ & \ddots & \\ & & h_K(Y_t;\theta) \end{bmatrix} V' \right)^{1/2} dW(t) \\
dW(t) \sim N(0, dtI)
\end{aligned}
\tag{7}
$$

#### S.2 Gaussian filtering and smoothing

We consider non-linear continuous-discrete state space models (CD-SSM) [1] of the form:

$$
\begin{aligned}
dx &= f(x,t,\theta)dt + L(x,t,\theta)dW \\
y_k &= g_k(x(t_k), r_k)
\end{aligned}
\tag{8}
$$

where $x(t) \in \mathbb{R}^n$ is the state at time $t$ and $y_k \in \mathbb{R}^d$ is the measurement collected at time $t_k$. The functions $f(x,t,\theta)$ and $g_k(x,r)$ define the dynamic and measurement models respectively, where $\{W(t) : t \geq 0\}$ is a $s$-dimensional Brownian motion with diffusion matrix $Q(t)$ and $\{r_k : k = 1,\ldots,K\}$ is multivariate Gaussian distributed white random sequence $r_k \sim \mathcal{N}_d(0, R_k)$ with (noise) covariance matrix $R_k$. We assume that the processes $\{W(t)\}_t$ and $\{r_k\}_k$ and the random initial condition $x(0) \sim N_n(m_0, P_0)$ are mutually independent. The matrix-valued function

$$
\Sigma(x,t,\theta) = L(x,t,\theta)Q(t)L'(x,t,\theta) \tag{9}
$$

defines the diffusion of the dynamic process, and it is state-dependent. In continuous-discrete filtering, the purpose is to compute the following filtering distributions

$$
p(x(t)|y_1,\ldots,y_k) \qquad t \in [t_k, t_{k+1}) \qquad k = 1,2,\ldots \tag{10}
$$

which is actually almost the same as the discrete filter where only the prediction step is replaced with solving of linearly-approximated Fokker–Planck–Kolmogorov (FPK) partial differential equations [2]. In continuous-discrete smoothing the goal is to estimate the following smoothing distributions

$$
p(x(t)|y_1,\ldots,y_K) \qquad t \in [t_0, t_K] \qquad k = 1,2,\ldots \tag{11}
$$

In this paper we focus on Gaussian filtering and smoothing [2, 3] where the distributions mentioned above are approximated as

$$\begin{aligned} p(x_k|y_{1:k-1},\theta) &= \mathcal{N}_n(m_k^*, P_k^*) \\ p(x_k|y_{1:k},\theta) &= \mathcal{N}_n(m_k, P_k) \\ p(y_k|y_{1:k-1},\theta) &= \mathcal{N}_d(\mu_k, S_k) \\ p(x_k|y_{1:K},\theta) &= \mathcal{N}_n\left(m_{k|K}^s, P_{k|K}^s\right) \end{aligned} \tag{12}$$

where $m_k$, $P_k$, $m_k^*$, $P_k^*$, $\mu_k$ and $S_k$ depend on $\theta$. These distributions are computed according to a step-wise algorithm defined below.

**1. Prediction step**: For every $k = 1, \ldots, K$ we consider the first-order continuous-discrete EKF prediction equations [1, 2]

$$\begin{cases} \frac{dm_k^*(t)}{dt} = f(m_k^*(t), t, \theta) \\ m_k^*(t_{k-1}) = m_{k-1} \end{cases} \tag{13a}$$

$$\begin{cases} \frac{dP_k^*(t)}{dt} = f_x(m_k^*(t), t, \theta)P_k^*(t) + P_k^*(t)f_x'(m_k^*(t), t, \theta) + \Sigma(m_k^*(t), t, \theta) \\ P_k^*(t_{k-1}) = P_{k-1} \end{cases} \tag{13b}$$

where $(\cdot)_x$ denotes the Jacobian operator w.r.t. $x$.

**2. Update step**: The solutions $m_k^*(t)$ and $P_k^*(t)$ obtained from the prediction step are used to update the next initial conditions via the equations

$$\begin{aligned} \mu_k &= g_k(m_k^*, r_k) \\ S_k &= g_{kx}(m_k^*, r_k)P_k^* g_{kx}'(m_k^*, r_k) + R_k \\ K_k &= P_k^* g_{kx}'(m_k^*, r_k)S_k^{-1} \\ m_k &= m_k^* + K_k(y_k - \mu_k) \\ P_k &= P_k^* - K_k S_k K_k' \end{aligned} \tag{14}$$

From these results, the marginal likelihood of the measurements given $\theta$ is given by [4–6]

$$p(y_1, \ldots, y_K|\theta) = \prod_{k=1}^{K} \mathcal{N}(y_k|\mu_k(\theta), S_k(\theta)) \tag{15}$$

where we highlight that the first two-order moments $\mu_k$'s and $S_k$'s depend both on $\theta$. Then, under the assumption of uniform prior for $\theta$, we update the parameters by minimizing the energy function $\varphi(\theta)$ [7] trough the following

**3. Optimization step**:

$$\theta \leftarrow \underset{\theta}{\texttt{argmin}} \underbrace{\sum_{k=1}^{K} \frac{1}{2} ln|2\pi S_k| + \frac{1}{2}\sum_{k=1}^{K}(y_k - \mu_k)' S_k^{-1}(y_k - \mu_k)}_{\varphi(\theta)} \tag{16}$$

which is effectively a maximum-likelihood step. The new parameter vector $\theta$ is then used to estimate the first two-order moments of the smoothing distribution $p(x_k|y_{1:K},\theta)$ using a backward step defined as follows

**4. Smoothing step**: We use the Rauch-Tung-Striebel Smoothing algorithm

(RTS) [8–10] and we estimate the first two-order moments of $p(x_k|y_{1:K},\theta)$ as

$$\begin{cases} B_{k+1} &= P_k(\theta)e^{f_x(m_k^*,t,\theta)}(P_{k+1}^*(\theta))^{-1} \\ m_{k|K}^s &= m_k(\theta) + B_{k+1}(m_{k+1|K}^s - m_{k+1}^*(\theta)) \\ P_{k|K}^s &= P_k(\theta) + B_{k+1}(P_{k+1|K}^s - P_{k+1}^*(\theta)) \end{cases} \tag{17}$$

where $e^{(\cdot)}$ is the matrix exponential operator and the values of $m_k$, $P_k$, $m_k^*$, $P_k^*$ are the ones already obtained from the filtering (prediction and update) steps. The first two terms of $B_{k+1}$ are obtained by solving the following initial value problem [2, 11]:

$$\begin{cases} \frac{dC_k^*(t)}{dt} &= C_k^*(t)f_x(m_k^*,t,\theta) \\ C_k^*(t_{k-1}) &= P_{k-1} \end{cases} \tag{18}$$

whose solution is $C_k^*(\theta) = P_{k-1}(\theta)e^{f_x(m_k^*,t,\theta)}$ in case $f_x(m_k^*,t,\theta)$ does not depend on $t$. Finally, we update the first two-order moments $m_0$ and $P_0$ of the initial condition $x_0$ as

**5. Update initial conditions**:

$$\begin{cases} m_0 \leftarrow m_{0|K}^s \\ P_0 \leftarrow P_{0|K}^s \end{cases} \tag{19}$$

As a summary, steps 1 - 2 define the Extended Kalman Filter (EKF). Step 4 defines the Rauch-Tung-Striebel Smoother (RTS). The 3rd step is needed to update the unknown parameters between every EKF and RTS steps. The last fifth step updates the mean and the covariance of the initial condition $x_0$. The whole EKF/RTS algorithm consists in running steps 1 - 5 until a convergence criterion is met for the free parameters $\theta$. What still is missing is to estimate the gradient $\nabla_\theta\varphi(\theta)$ of the energy function $\varphi(\theta)$. This requires $p$ more prediction steps and $p$ more update steps, each for every component of $\theta$. We show these additional steps only in next section for the filtering/smoothing problem of a state space model having as dynamic component a stochastic quasi-reaction network, for which most of the previous formulations simplify.

##### S.2.1 Prediction/Update derivatives

We consider the continuous-discrete state space model (2)-(3) from the main text. The computation of the gradient $\nabla_{\theta,\rho_0,\rho_1}\varphi(\theta,\rho_0,\rho_1)$ of the energy function $\varphi(\theta,\rho_0,\rho_1)$ requires, at every time point $k$, $p+2$ more prediction and update steps in order to compute the terms $\frac{\partial S_k}{\partial\theta_j}$'s, $\frac{\partial\mu_k}{\partial\theta_j}$'s, $\frac{\partial S_k}{\partial\rho_0}$, $\frac{\partial\mu_k}{\partial\rho_0}$, $\frac{\partial S_k}{\partial\rho_1}$ and $\frac{\partial\mu_k}{\partial\rho_1}$, where $p$ is the dimension of $\theta$. These are obtained by deriving the equations in (4) and (6) from the main text w.r.t. $\theta$, $\rho_0$ and $\rho_1$ as shown below.

**1'. Prediction derivatives step**:

"$\boldsymbol{\frac{\partial m_k^*}{\partial\theta_j}}$": If we derive the system (4a) from the main text w.r.t. $\theta$ we get

$$\begin{cases} \frac{d}{dt}\left(\frac{\partial}{\partial\theta_j}m_k^*(t)\right) = \frac{\partial}{\partial\theta_j}V_\theta m_k^*(t) = V_\theta\frac{\partial}{\partial\theta_j}m_k^*(t) + \left(\frac{\partial}{\partial\theta_j}V_\theta\right)m_k^*(t) \\ \frac{\partial}{\partial\theta_j}m_k^*(t_{k-1}) = \frac{\partial}{\partial\theta_j}m_{k-1} \end{cases}$$

By using the integrating factor $I = e^{-\int_{t_{k-1}}^{t} V_\theta ds} = e^{-V_\theta(t-t_{k-1})}$ we get

$$\frac{\partial m_k^*}{\partial \theta_j} = e^{V_\theta(t-t_{k-1})} \left\{ \int_{t_{k-1}}^{t} e^{-V_\theta(s-t_{k-1})} \left( \frac{\partial}{\partial \theta_j} V_\theta \right) m_k^*(s) ds + \frac{\partial}{\partial \theta_j} m_{k-1} \right\}$$

$$\Rightarrow \frac{\partial m_k^*}{\partial \theta_j} = \int_{t_{k-1}}^{t} e^{V_\theta(t-s)} \frac{\partial}{\partial \theta_j} V_\theta e^{V_\theta(s-t_{k-1})} m_{k-1} ds + e^{V_\theta(t-t_{k-1})} \frac{\partial}{\partial \theta_j} m_{k-1} \quad (20)$$

"$\boldsymbol{\frac{\partial P_k^*}{\partial \theta_j}}$": By deriving the system (4b) from the main text w.r.t. $\theta$ we get

$$\begin{aligned} \frac{d}{dt}\left( \frac{\partial}{\partial \theta_j} P_k^*(t) \right) &= \frac{\partial}{\partial \theta_j} \left\{ V_\theta P_k^*(t) + P_k^*(t) V_\theta' + \Delta t \beta(m_k^*(t), \theta) \right\} \\ &= \frac{\partial}{\partial \theta_j} V_\theta P_k^*(t) + V_\theta \frac{\partial}{\partial \theta_j} P_k^*(t) + \frac{\partial}{\partial \theta_j} P_k^*(t) V_\theta' + P_k^*(t) \frac{\partial}{\partial \theta_j} V_\theta' + \\ &+ \Delta t \left\{ \sum_{i=1}^{n} \frac{\partial \beta(m_k^*(t), \theta)}{\partial x_i} \frac{\partial m_{ki}^*(t)}{\partial \theta_j} + \frac{\partial \beta(m_k^*(t), \theta)}{\partial \theta_j} \right\} \\ &= V_\theta \frac{\partial}{\partial \theta_j} P_k^*(t) + \frac{\partial}{\partial \theta_j} P_k^*(t) V_\theta' + Q(t) \end{aligned}$$

where

$$Q(t) = \frac{\partial}{\partial \theta_j} V_\theta P_k^*(t) + P_k^*(t) \frac{\partial}{\partial \theta_j} V_\theta' + \Delta t \left\{ \sum_{i=1}^{n} \frac{\partial \beta(m_k^*(t), \theta)}{\partial x_i} \frac{\partial m_{ki}^*(t)}{\partial \theta_j} + \frac{\partial \beta(m_k^*(t), \theta)}{\partial \theta_j} \right\}$$

which is a differential Sylvester equation [12]. The corresponding initial value problem is

$$\begin{cases} \frac{d}{dt}\left( \frac{\partial}{\partial \theta_j} P_k^*(t) \right) = V_\theta \frac{\partial}{\partial \theta_j} P_k^*(t) + \frac{\partial}{\partial \theta_j} P_k^*(t) V_\theta' + Q(t) \\ \frac{\partial}{\partial \theta_j} P_k^*(t_{k-1}) = \frac{\partial}{\partial \theta_j} P_{k-1} \end{cases}$$

whose solution is given by [12]

$$\frac{\partial}{\partial \theta_j} P_k^*(t) = e^{(t-t_{k-1})V_\theta} \frac{\partial}{\partial \theta_j} P_{k-1} e^{(t-t_{k-1})V_\theta'} + \int_{t_{k-1}}^{t} e^{(t-s)V_\theta} Q(s) e^{(t-s)V_\theta'} ds \quad (21)$$

"$\boldsymbol{\frac{\partial m_k^*}{\partial \rho_0}}$": After deriving the system (4a) from the main text w.r.t. $\rho_0$ we get

$$\begin{cases} \frac{d}{dt}\left( \frac{\partial}{\partial \rho_0} m_k^*(t) \right) = V_\theta \frac{\partial}{\partial \rho_0} m_k^*(t) \\ \frac{\partial}{\partial \rho_0} m_k^*(t_{k-1}) = \frac{\partial}{\partial \rho_0} m_{k-1} \end{cases}$$

By using the integrating factor $I = e^{-V_\theta(t-t_{k-1})}$ we get the following solution

$$\frac{\partial}{\partial \rho_0} m_k^*(t) = e^{V_\theta(t-t_{k-1})} \frac{\partial}{\partial \rho_0} m_{k-1} \quad (22)$$

"$\frac{\partial P_k^*}{\partial \rho_0}$": Finally we derive the system (4b) from the main text w.r.t. $\rho_0$ and we get

$$\begin{cases} \frac{d}{dt}\left(\frac{\partial}{\partial \rho_0} P_k^*(t)\right) = V_\theta \frac{\partial}{\partial \rho_0} P_k^*(t) + \frac{\partial}{\partial \rho_0} P_k^*(t) V_\theta' + \Delta t \sum_{i=1}^n \frac{\partial \beta(m_k^*(t),\theta)}{\partial x_i} \frac{\partial m_{ki}^*(t)}{\partial \rho_0} \\ \frac{\partial}{\partial \rho_0} P_k^*(t_{k-1}) = \frac{\partial}{\partial \rho_0} P_{k-1} \end{cases}$$

which is a differential Sylvester initial value problem whose solution is given by [12]

$$\frac{\partial}{\partial \rho_0} P_k^*(t) = e^{(t-t_{k-1})V_\theta} \frac{\partial}{\partial \rho_0} P_{k-1} e^{(t-t_{k-1})V_\theta'} + \int_{t_{k-1}}^t e^{(t-s)V_\theta} Q(s) e^{(t-s)V_\theta'} ds \quad (23)$$

where

$$Q(t) = \Delta t \sum_{i=1}^n \frac{\partial \beta(m_k^*(t),\theta)}{\partial x_i} \frac{\partial m_{ki}^*(t)}{\partial \rho_0}$$

The formulations of $\frac{\partial m_k^*}{\partial \rho_1}$ and $\frac{\partial P_k^*}{\partial \rho_1}$ are equivalent to the case of $\rho_0$. The resulting solutions $\frac{\partial m_k^*}{\partial \theta_j}$, $\frac{\partial P_k^*}{\partial \theta_j}$, $\frac{\partial m_k^*}{\partial \rho_0}$, $\frac{\partial P_k^*}{\partial \rho_0}$, $\frac{\partial m_k^*}{\partial \rho_1}$ and $\frac{\partial P_k^*}{\partial \rho_1}$ are then used to update the corresponding initial values via a set of equations obtained by deriving all the equations of (6) from the main text w.r.t. $\psi = (\theta, \rho_0, \rho_1)$ as shown below.

**2'. Update derivatives step**:

$\theta$:

$$\begin{aligned} \frac{\partial \mu_k}{\partial \theta_j} &= G_k \frac{\partial m_k^*}{\partial \theta_j}; \qquad \frac{\partial S_k}{\partial \theta_j} = G_k \frac{\partial P_k^*}{\partial \theta_j} G_k' + R_k \\ \frac{\partial K_k}{\partial \theta_j} &= \frac{\partial P_k^*}{\partial \theta_j} G_k' S_k^{-1} - P_k^* G_k' S_k^{-1} \frac{\partial S_k}{\partial \theta_j} S_k^{-1} \\ \frac{\partial m_k}{\partial \theta_j} &= \frac{\partial m_k^*}{\partial \theta_j} + \frac{\partial K_k}{\partial \theta_j}(y_k - \mu_k) - K_k \frac{\partial \mu_k}{\partial \theta_j} \\ \frac{\partial P_k}{\partial \theta_j} &= \frac{\partial P_k^*}{\partial \theta_j} - \frac{\partial K_k}{\partial \theta_j} S_k K_k' - K_k \frac{\partial S_k}{\partial \theta_j} K_k' - K_k S_k \frac{\partial K_k}{\partial \theta_j}' \end{aligned} \quad (24)$$

$\rho_0$:

$$\begin{aligned} \frac{\partial \mu_k}{\partial \rho_0} &= G_k \frac{\partial m_k^*}{\partial \rho_0}; \qquad \frac{\partial S_k}{\partial \rho_0} = G_k \frac{\partial P_k^*}{\partial \rho_0} G_k' + I_d + \rho_1 diag\left(\frac{\partial \mu_k}{\partial \rho_0}\right) \\ \frac{\partial K_k}{\partial \rho_0} &= \frac{\partial P_k^*}{\partial \rho_0} G_k' S_k^{-1} - P_k^* G_k' S_k^{-1} \frac{\partial S_k}{\partial \rho_0} S_k^{-1} \\ \frac{\partial m_k}{\partial \rho_0} &= \frac{\partial m_k^*}{\partial \rho_0} + \frac{\partial K_k}{\partial \rho_0}(y_k - \mu_k) - K_k \frac{\partial \mu_k}{\partial \rho_0} \\ \frac{\partial P_k}{\partial \rho_0} &= \frac{\partial P_k^*}{\partial \rho_0} - \frac{\partial K_k}{\partial \rho_0} S_k K_k' - K_k \frac{\partial S_k}{\partial \rho_0} K_k' - K_k S_k \frac{\partial K_k}{\partial \rho_0}' \end{aligned} \quad (25)$$

$\rho_1$:

$$
\begin{aligned}
\frac{\partial \mu_k}{\partial \rho_1} &= G_k \frac{\partial m_k^*}{\partial \rho_1}; \qquad \frac{\partial S_k}{\partial \rho_1} = G_k \frac{\partial P_k^*}{\partial \rho_1} G_k' + diag(\mu_k) + \rho_1 diag\left(\frac{\partial \mu_k}{\partial \rho_1}\right) \\
\frac{\partial K_k}{\partial \rho_1} &= \frac{\partial P_k^*}{\partial \rho_1} G_k' S_k^{-1} - P_k^* G_k' S_k^{-1} \frac{\partial S_k}{\partial \rho_1} S_k^{-1} \\
\frac{\partial m_k}{\partial \rho_1} &= \frac{\partial m_k^*}{\partial \rho_1} + \frac{\partial K_k}{\partial \rho_1}(y_k - \mu_k) - K_k \frac{\partial \mu_k}{\partial \rho_1} \\
\frac{\partial P_k}{\partial \rho_1} &= \frac{\partial P_k^*}{\partial \rho_1} - \frac{\partial K_k}{\partial \rho_1} S_k K_k' - K_k \frac{\partial S_k}{\partial \rho_1} K_k' - K_k S_k \frac{\partial K_k}{\partial \rho_1}'
\end{aligned}
\tag{26}
$$

#### S.3 Simulation studies

Consider the cell differentiation network of Figure S.1 written in the state space model formulation

$$
\begin{aligned}
x_{t+1} - x_t = \Delta x &= V h(x_t;\theta)\Delta t + \underbrace{\left(V \begin{bmatrix} h_1(x_t;\theta) & & \\ & \ddots & \\ & & h_J(x_t;\theta) \end{bmatrix} V'\right)^{1/2}}_{\beta(x_t;\theta)} dW_t \\
y_k &= G_k x(t_k) + r_k
\end{aligned}
\tag{27}
$$

where

$$
dW_t \sim \mathcal{N}_n(0, \Delta t I_n); \quad r_k \sim \mathcal{N}_d(0, R_k); \quad R_k = \rho_0 I_d + \rho_1 diag(G_k x(t_k)) \tag{28}
$$

The net-effect matrix $V$ and an hazard vector $h(x,\theta)$ are defined as

$$
v_k = \begin{cases} (0 \dots 1_i \dots 0)' \\ (0 \dots -1_i \dots 0)' \\ (0 \dots -1_i \dots 2_j \dots 0)' \end{cases} \qquad h_k(X_t, \theta_i) = \begin{cases} X_{it}\alpha_i & \text{duplication} \\ X_{it}\delta_i & \text{death} \\ X_{it}\lambda_{ij} & \text{differentiation} \end{cases} \tag{29}
$$

where $i \neq j = 1, \dots, N$ and $\theta$ is the vector of the unknown parameters. We assume that the information on the HSCs and progenitors P1 and P2 is not available at every time point, and therefore we consider them as a latent states which cannot be measured. Furthermore, all the lineages that have not been recaptured for a particular clone at a given time point are also considered as latent states. Therefore, for the measurement model the selection matrix $G_k$ is defined accordingly. In our simulation studies we also assume the following conservation laws

$$
\begin{aligned}
\lambda_{HSC\to P1} &= \lambda_{P1\to T} + \lambda_{P1\to B} + \lambda_{P1\to NK} \\
\lambda_{HSC\to P2} &= \lambda_{P2\to G} + \lambda_{P2\to M}
\end{aligned}
\tag{30}
$$

so as to facilitate the inference of parameters related to the systematically missing cell types. In each simulation study, to generate the clone-specific trajectories, we use the Euler-Maruyama method [13] with an initial condition $x_0 = 100$ for the HSCs and zero otherwise. Each trajectory, starting from $t_0 = 0$ and terminating at $t_1 = 1$, has a sample size equal to 1000 with $\Delta t = 1/1000$. Then, we select a subset of $T$ equidistant time points, where $T$ is chosen depending on the

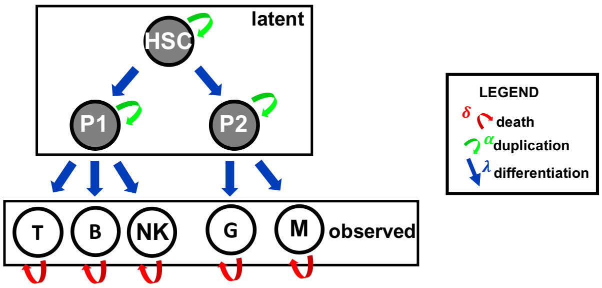


Figure S.1: A cell differentiation network of eight cell types (nodes). The information on the HSC and the progenitors P1 and P2 are missing. Red arrows denote a death move, green arrows indicate a duplication move, and blue arrows a differentiation move.

particular simulation design. Each simulation study is designed to test parameter uncertainty when reducing $T$ (figure S.2), reducing the fraction $0 < f < 1$ of clones recaptured across lineages and time (figure S.3), increasing measurement noise parameters $(\rho_0, \rho_1)$ (figure S.4), selecting a cell differentiation structure among the candidates (figure S.5).

Results from simulations show accurate performance of the method in the identification of the missing states and in the inference of the true parameters. In particular, results from simulation 1 suggest that reducing the number of time points of the complete simulated trajectories does not affect parameter inference, even in the extreme case where we fitted our model to the data of only five time points out of 1000 of the complete trajectories. Second simulation clearly indicate that our method still provides good estimates if we reduce the fraction $f$ of the observed states, even for an high fraction of missing states ($f = 0.1$). Robustness on measurement noise has been also assessed in simulation 3, whose results show that parameters are identifiable, even under extreme noise setting ($\rho_0 = \rho_1 = 100$) where still we get some sensible estimates. Finally, in the fourth simulation study our method combined with Akaike Information Criterion was able to select the correct structure among the candidates. Furthermore, we empirically proved that considering missing values as latent states is of crucial importance. To this end we simulated an additional synthetic clonal tracking dataset and we randomly set 50% of the simulated trajectories to missing/not available. Then we first fitted our model with the missing states set to zero, and subsequently we fitted the same model considering the missing states as latent states. Results are shown in Figure S.6 and clearly indicate that considering missing values as low counts (e.g. zero) only provides a poor estimate for both the parameters and the smoothing moments. Whereas, if we treat missing counts as latent states, the parameter estimates are near the true ones and the smoothing moments correctly fit the stochastic process.

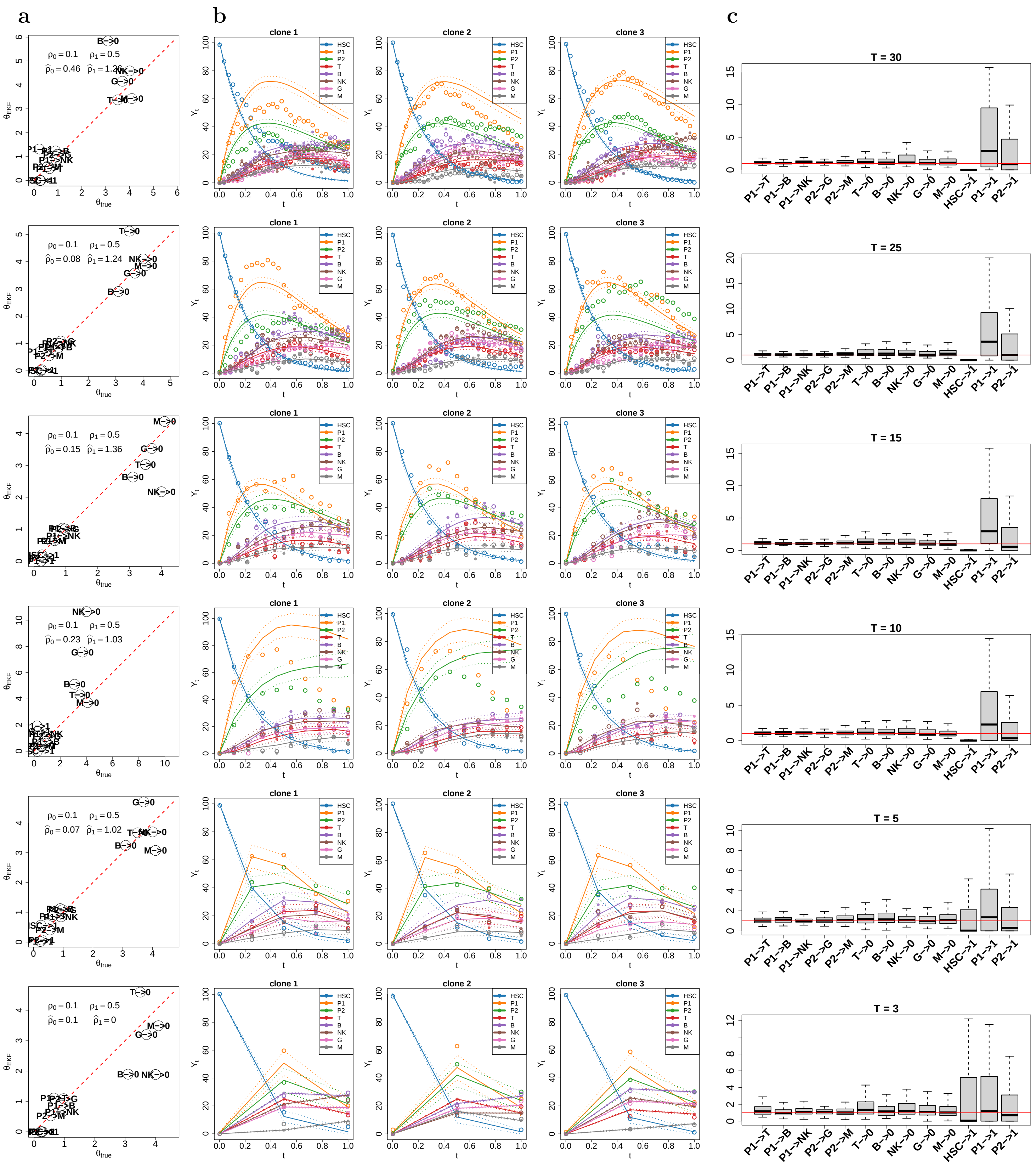


Figure S.2: For each $T$ (rows): Scatterplot of the estimated parameters $\hat{\theta}_{EKF}$ against the true parameters $\theta_{true}$ (a). The simulated process $\{x_t\}_t$ (empty dots), the noise-corrupted measurements $\{y_k\}_k$ (full dots), and the estimated smoothing moments $m^s_{k|K}$ and $P^s_{k|K}$ for the three clones (b). Histograms of the estimated parameters over 100 independent simulations (c).

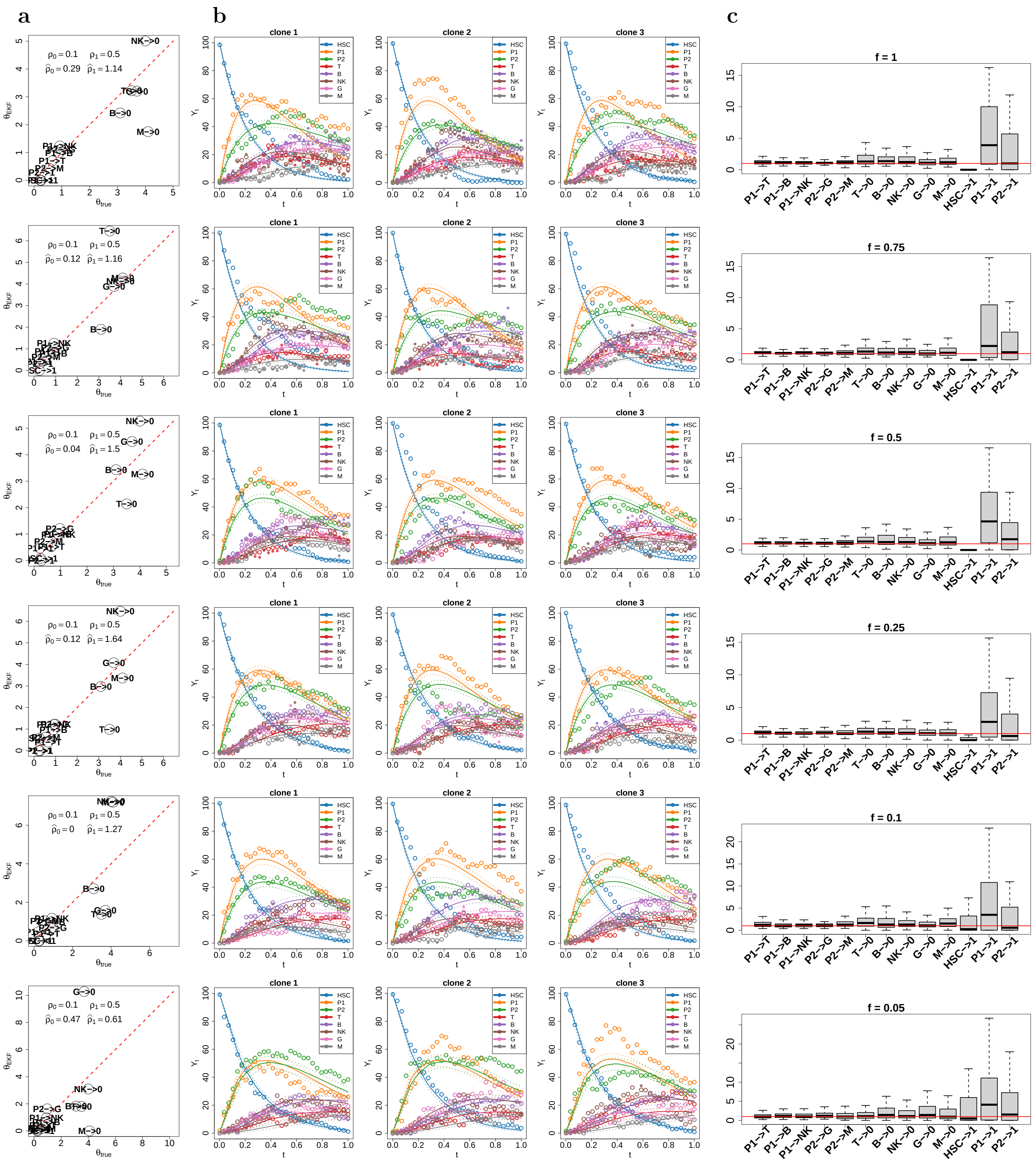


Figure S.3: For each $f$ (rows): Scatterplot of the estimated parameters $\hat{\theta}_{EKF}$ against the true parameters $\theta_{true}$ (a). The simulated process $\{x_t\}_t$ (empty dots), the noise-corrupted measurements $\{y_k\}_k$ (full dots), and the estimated smoothing moments $m^s_{k|K}$ and $P^s_{k|K}$ for the three clones (b). Histogram of the estimated process parameters over 100 independent simulations (c).

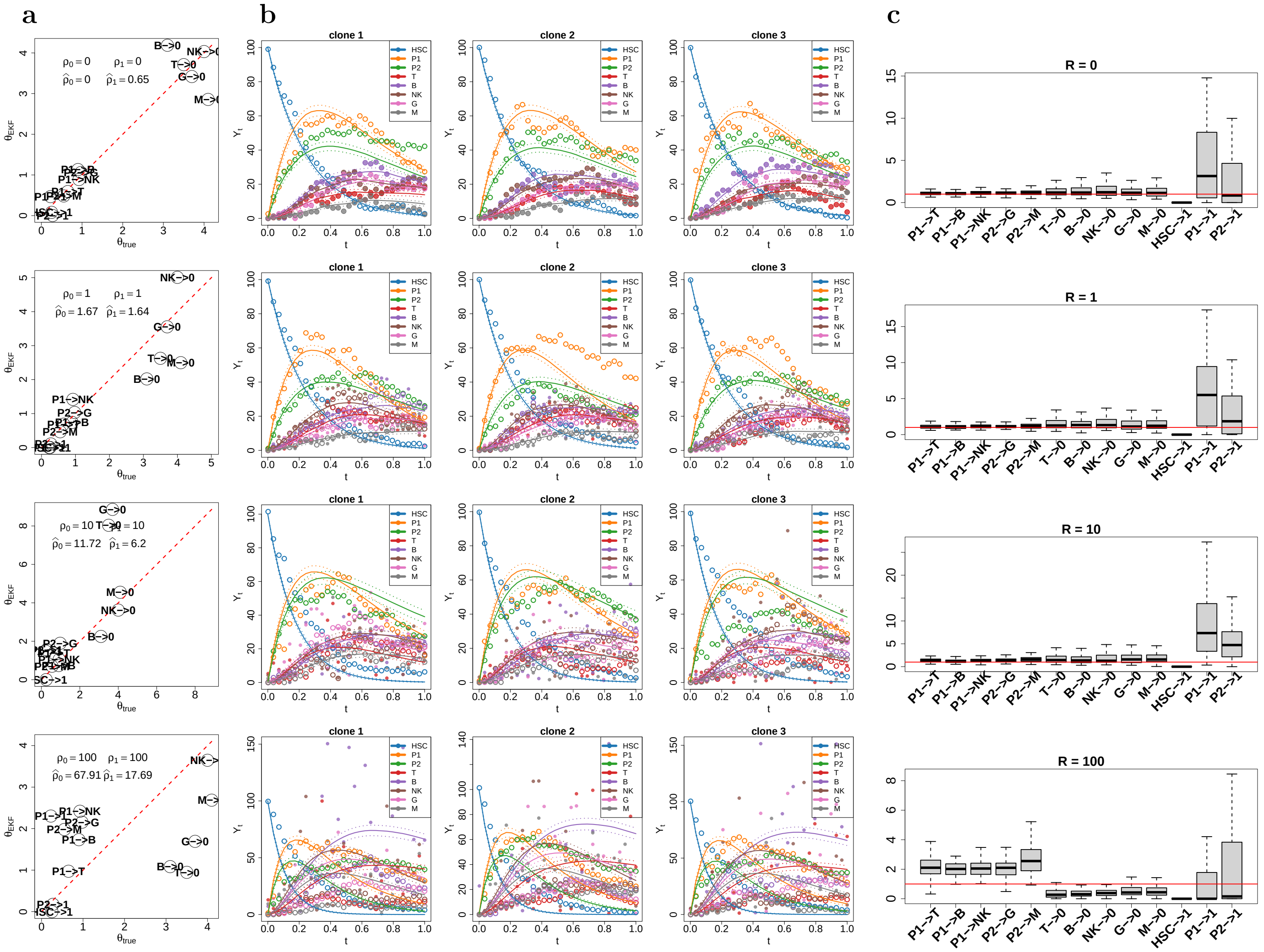

Figure S.4: For each $\rho_0, \rho_1 = 0, 1, 10, 100$ (rows): Scatterplot of the estimated parameters $\hat{\theta}_{EKF}$ against the true parameters $\theta_{true}$ (a). The simulated process $\{x_t\}_t$ (empty dots), the noise-corrupted measurements $\{y_k\}_k$ (full dots), and the estimated smoothing moments $m^s_{k|K}$ and $P^s_{k|K}$ for the three clones (b). Histogram of the estimated process parameters over 100 independent simulations (c).

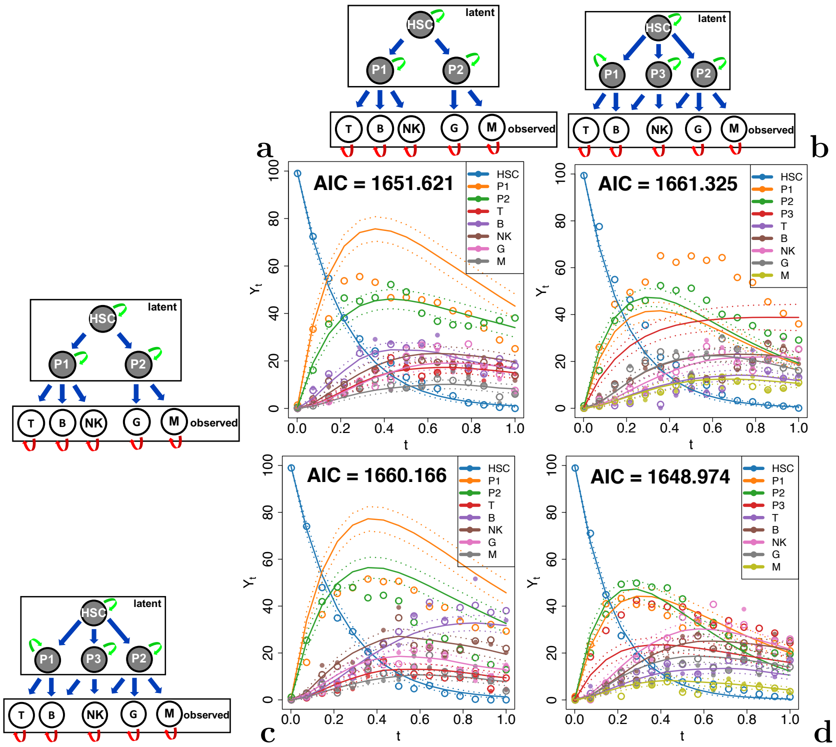

Figure S.5: For each true model (row) and for each candidate model (column): The simulated process $\{x_t\}_t$ (empty dots), the noise-corrupted measurements $\{y_k\}_k$ (full dots), and the estimated smoothing moments $m^s_{k|K}$ and $P^s_{k|K}$. The median AIC across all simulations is reported at the top of each plot panel. The smoothing moments of only one clone are shown in each case.

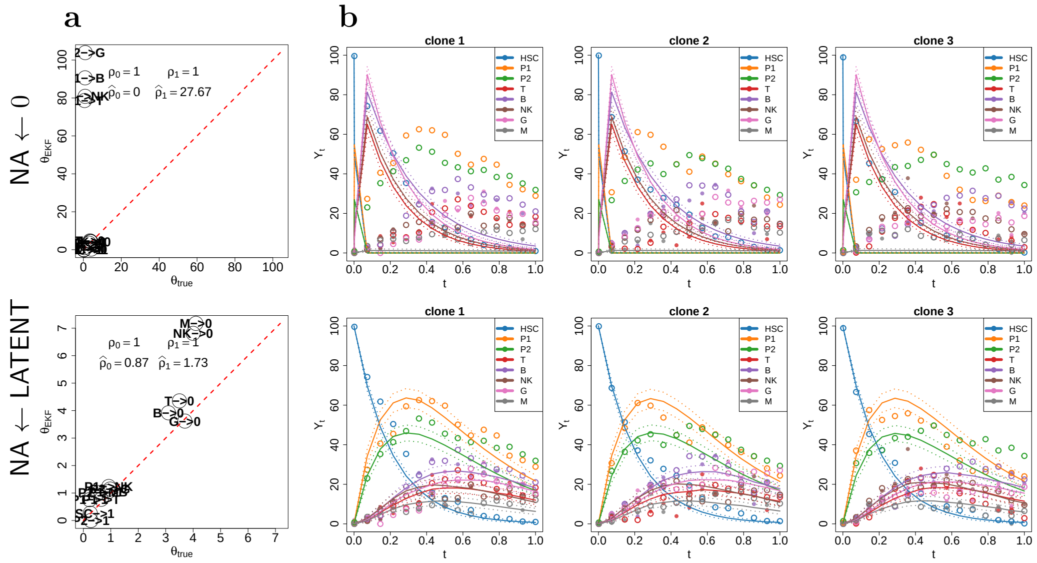

Figure S.6: NA $\leftarrow$ 0 (top) and NA $\leftarrow$ LATENT bottom: Scatterplot of the estimated parameters $\hat{\theta}_{EKF}$ against the true parameters $\theta_{true}$, along with the true and estimated noise parameters in the top-left corner (a). Values for the simulated process $\{x_t\}_t$ (empty dots), the noise-corrupted measurements $\{y_k\}_k$ (full dots), and the estimated smoothing moments $m^s_{k|K}$ and $P^s_{k|K}$ for the three clones (b). In this case only one simulation has been performed.

#### S.4 Karen pseudocode

**Input:** $\{x_k\}_k$, $V$, $h(x,\theta)$, $x_0 \sim \mathcal{N}_n(m_0, P_0)$
**Output:** $\hat{\theta}_{ekf}$, $\hat{\rho}_{0ekf}$, $\hat{\rho}_{1ekf}$, $m^s_{k|K}$ and $P^s_{k|K}$
**while** $\epsilon > \texttt{tol}$ **do**
  $\theta_{old} \leftarrow \theta$
  **for** $k = 1 : K$ **do**
    **1. Prediction:** `get` $m^*_k$ `and` $P^*_k$
    **1'. Prediction derivatives:** `get` $\frac{\partial m^*_k}{\partial \theta_j}$, $\frac{\partial}{\partial \theta_j} P^*_k$, $\frac{\partial m^*_k}{\partial \rho_0}$, $\frac{\partial}{\partial \rho_0} P^*_k$, $\frac{\partial m^*_k}{\partial \rho_1}$ `and` $\frac{\partial}{\partial \rho_1} P^*_k$
    **2. Update:** `get` $m_k$, $P_k$, $\mu_k$ `and` $S_k$
    **2'. Update derivatives:** `get` $\frac{\partial m_k}{\partial \theta_j}$, $\frac{\partial}{\partial \theta_j} P_k$, $\frac{\partial m_k}{\partial \sigma^2}$, $\frac{\partial \mu_k}{\partial \theta_j}$, $\frac{\partial}{\partial \theta_j} S_k$, $\frac{\partial}{\partial \rho_0} P_k$, $\frac{\partial \mu_k}{\partial \rho_0}$, $\frac{\partial}{\partial \rho_0} S_k$, $\frac{\partial}{\partial \rho_1} P_k$, $\frac{\partial \mu_k}{\partial \rho_1}$, $\frac{\partial}{\partial \rho_1} S_k$
  **end**
  **3. Optimization:**

$$(\theta, \rho_0, \rho_1) \leftarrow \underset{\theta \geq 0, \rho_0 > 0, \rho_1 > 0}{\texttt{argmin}} \underbrace{\sum_{k=1}^{K} \frac{1}{2} ln|2\pi S_k| + \frac{1}{2} \sum_{k=1}^{K} (y_k - \mu_k)' S_k^{-1} (y_k - \mu_k)}_{\varphi(\theta, \rho_0, \rho_1)}$$

  **4. Smoothing:** `Get` $m^s_{k|K}$ `and` $P^s_{k|K}$
  **5. Update prior:** `Update prior distribution for` $x_0$ `as`

$$m_0 \leftarrow m^s_{1|K}$$
$$P_0 \leftarrow P^s_{1|K}$$

  $\epsilon \leftarrow \frac{\|\theta - \theta_{old}\|_2}{\|\theta_{old}\|_2}$
**end**

**Algorithm 1:** Pseudocode of Karen. The analytical formulations of the prediction/update steps for the filtering/smoothing moments, the marginal likelihood, and the corresponding derivatives, can be found in equations (4)-(10) from Online Methods.

#### S.5 Rhesus macaque data rescaling

Although the sample DNA amount was maintained constant during the whole experiment (200 ng for ZH33 and ZG66 or 500 ng for ZH17), the sample collected resulted in different magnitudes of total number of reads. Table S.1 shows the total number of reads collected in each sample of the rhesus macaque clonal tracking dataset. This discrepancy makes all the samples not comparable across time and cell types. Therefore we rescaled the barcode counts according to

$$Y_{ijk} \leftarrow Y_{ijk} \cdot \frac{\min_{ij} \sum_k Y_{ij}}{\sum_k Y_{ij}} \tag{31}$$

where $Y_{ijk}$ is the $ijk$-entry of the barcode matrix with dimensions $(i, j, k)$ mapping respectively time, cell type and clone.

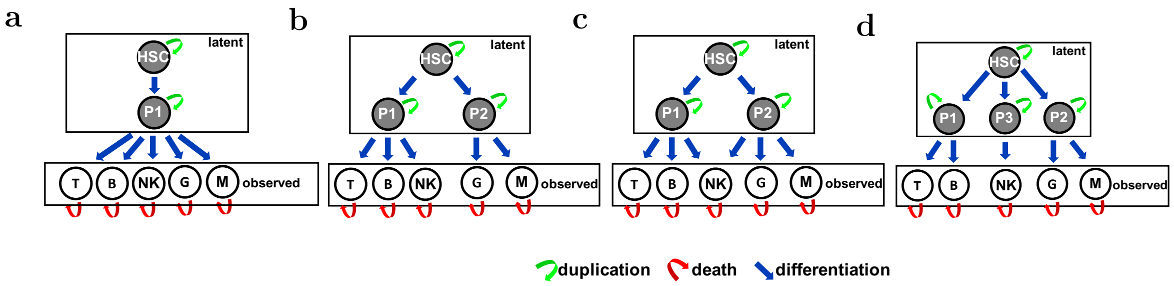


Figure S.7: Graphical representation of the candidate models: Latent and observed cell types are indicated with grey and white nodes respectively. Red arrows denote a death move, green arrows indicate a duplication move, and blue arrows a differentiation move.

| | | T | B | NK | M | G |
|---|---|---|---|---|---|---|
| ZH33 | 1 | 1465289 | 74735 | 135092 | 119331 | 2831 |
| | 2 | 225797 | 216844 | 335789 | 1035270 | 908685 |
| | 3 | 243986 | 413757 | 663184 | 886682 | 816990 |
| | 4.5 | 485542 | 479493 | 834064 | 985821 | 987171 |
| | 6.5 | 645005 | 676413 | 926089 | 895309 | 911637 |
| | 9.5 | 829073 | 962325 | 1057398 | 1229233 | 1220506 |
| ZH17 | 1 | 51802 | 1347050 | 1288718 | 1351450 | 707382 |
| | 2 | 826190 | 1342700 | 1350703 | 1354355 | 1213749 |
| | 3 | 1303922 | 1347692 | 1338024 | 1347177 | 1283250 |
| | 4.5 | 190591 | 1206361 | 489098 | 572877 | 1195585 |
| | 6.5 | 887851 | 610999 | 1344488 | 381552 | 1339299 |
| ZG66 | 1 | 752127 | 0 | 211350 | 13382 | 0 |
| | 2 | 692133 | 58890 | 308800 | 363310 | 145252 |
| | 3 | 339292 | 209137 | 424458 | 808404 | 704331 |
| | 4.5 | 617281 | 338977 | 718472 | 887183 | 897672 |

Table S.1: Total number of reads (sum across the different clones) collected in each treated animal at each time point and for all the cell types.

#### S.6 Hematopoietic models

In this work we consider four different biologically-sustained models of hematopoiesis whose graphical representation is shown in Figure S.7. Model (A) is a single-branch developmental tree where the hematopoietic stem cells produces all the mature cell type trough a single multipotent intermediate progenitor $P_1$. According to model (B) the lymphoid cells (T, B, NK) and the myeloid cells (G, M) are generated trough separate branches of differentiation. Therefore, this is very similar to the well known classical/dichotomic model of hematopoiesis [14]. This model classifies blood cells into two major lineages, but finally differentiated cells are placed in parallel. In contrast, model (C) proposes the idea that myeloid cells represent a prototype of hematopoietic cells capable to produce both myeloid G/M cells and lymphoid NK cells, whereas T, B and NK cells represent specialized types. Therefore model C can be interpreted as the myeloidbased model from [14]. Finally, model (D) assumes that while lymphoid T/B and myeloid G/M develop in parallel trough separate branches from different progenitors, there is a third developmental branch for the NK cells which is separated/independent from the first two branches.

#### S.7 Karen R package: minimal working examples

In this section we describe some key functionalities of our R package Karen. Section S.7.1 shows how to simulate a clonal tracking dataset from a stochastic quasi-reaction network defined by a given set of biochemical reactions. In particular, we show how to simulate clone-specific stochastic trajectories. Section S.7.2 shows how to fit a Kalman Reaction Network to a simulated clonal tracking dataset. Finally in Section S.7.3 we analyse a simulated clonal tracking dataset with Karen and we show how to visualize the results.

##### S.7.1 Simulating clonal tracking datasets

A clonal tracking dataset compatible with Karen's functions must be formatted as a 3-dimensional array $Y$ whose $ijk$-entry $Y_{ijk}$ is the number of cells of clone $k$ for cell type $j$ collected at time $i$. The function `get.sim.trajectories()` can be used to simulate clone-specific trajectories given an initial condition $X_0$ for a set of observed `ct.lst` and latent `latSts.lst`, and obeying to a particular cell differentiation network defined by a list `rct.lst` of biochemical reactions, subject to a set of linear constraints `constr.lst`. The cellular events of duplication, death and differentiation are respectively coded in the package with the character labels `"A->1"`, `"A->0"`, and `"A->B"`, where `A` and `B` are two distinct cell types. The cell types provided in $Y$, `ct.lst`, `latSts.lst`, `rct.lst` and `constr.lst` must be compatible, otherwise an error will be raised (see Karen documentation for details).

The following R code chunk shows how to simulate clone-specific trajectories of cells using the Euler-Maruyama method [13]. As an illustrative example we focus on the cell differentiation network structure from Figure S.7-b having eight synthetic cell types. Here we assume that the hematopoietic stem cells HSC, and the two intermediate progenitors P1 - P2 are latent cell types that cannot be measured. We also assume a recapture rate $f_{NA} = .75$ and $(\rho_0 = .1, \rho_1 = .5)$ as the measurement noise parameters (see Section S.3 for details).

```
cat("\nInstall/load packages")
inst.pkgs <- installed.packages() ## installed packages

## required packages:
l.pkgs <- c("expm",
            "Matrix",
            "parallel",
            "gaussquad",
            "splines",
            "scales",
            "mvtnorm",
            "tmvtnorm",
            "MASS",
            "igraph",
            "stringr",
```

```
             "devtools")
## check if packages are installed:
lapply(l.pkgs, function(pkg){
  if(!(pkg %in% rownames(inst.pkgs))){
    install.packages(pkg)
  }
})
## load packages:
lapply(l.pkgs, function(pkg){library(pkg,character.only=TRUE
    ↪ )})

## install and load Karen:
if(!("Karen" %in% rownames(inst.pkgs))){
  install_github("delcore-luca/Karen",
                 ref = "master")
}
library(Karen)

rm(list = ls())

rcts <- c("HSC->P1", ## reactions
          "HSC->P2",
          "P1->T",
          "P1->B",
          "P1->NK",
          "P2->G",
          "P2->M",
          "T->0",
          "B->0",
          "NK->0",
          "G->0",
          "M->0"
          ,"HSC->1"
          ,"P1->1"
          ,"P2->1"
)

cnstr <- c("theta\\[\\'HSC->P1\\'\\]=(theta\\[\\'P1->T\\'\\]
    ↪  + theta\\[\\'P1->B\\'\\] + theta\\[\\'P1->NK\\'\\])",
           "theta\\[\\'HSC->P2\\'\\]=(theta\\[\\'P2->G\\'\\]
               ↪  + theta\\[\\'P2->M\\'\\])") ## reaction
               ↪ constraints
latsts <- c("HSC", "P1", "P2") ## latent cell types

ctps <- unique(setdiff(c(sapply(rcts, function(r){ ## all
    ↪ cell types
  as.vector(unlist(strsplit(r, split = "->", fixed = T)))
}, simplify = "array")), c("0", "1")))


########## TRUE PARAMETERS ##########
th.true <- c(0.65, 0.9, 0.925, 0.975, 0.55, 3.5, 3.1, 4,
    ↪ 3.7, 4.1, 0.25, 0.225, 0.275) ## dynamic parameters
names(th.true) <- tail(rcts, -length(cnstr))
```

```
s2.true <- 1e-8 ## additonal noise
r0.true <- .1 ## intercept noise parameter
r1.true <- .5 ## slope noise parameter
phi.true <- c(th.true, r0.true, r1.true) ## whole vector
    ↪ parameter
names(phi.true) <- c(names(th.true), "r0", "r1")

########## SIMULATION PARAMETERS ##########
S <- 1000 ## trajectories length
nCL <- 3 ## number of clones
X0 <- rep(0, length(ctps)) ## initial condition
names(X0) <- ctps
X0["HSC"] <- 100
ntps <- 30 ## number of time-points
f_NA <- .75 ## fraction of observed data

########## SIMULATE TRAJECTORIES ##########
XY <- get.sim.trajectories(rct.lst = rcts,
                           constr.lst = cnstr,
                           latSts.lst = latsts,
                           ct.lst = ctps,
                           th = th.true,
                           S = S,
                           nCL = nCL,
                           X0 = X0,
                           s2 = s2.true,
                           r0 = r0.true,
                           r1 = r1.true,
                           f = f_NA,
                           ntps = ntps,
                           trunc = FALSE)

XY$X ## process
XY$Y ## measurements
```

##### S.7.2 Fitting a Kalman Reaction Network

In this section we show how to fit a Kalman Reaction Network to a clonal tracking dataset. To this end we use the function `get.fit()` that includes the inference procedure described in the Online Methods section from the main paper. This function receives several arguments as input, such as the first two-order moments $m_0$ and $P_0$ of the initial condition $X_0$, a set of observed `ct.lst` and latent `latSts.lst` following a particular cell differentiation structure defined by a list `rct.lst` of biochemical reactions, subject to a set of linear constraints `constr.lst`. The function receives also a list of optimization parameters (see Karen documentation for details). As for the `get.sim.trajectories()` function, the cellular events of duplication, death and differentiation are respectively coded in the package with the character labels `"A->1"`, `"A->0"`, and `"A->B"`, where `A` and `B` are two distinct cell types. The cell types provided in $Y$, $m_0$, `ct.lst`, `latSts.lst`, `rct.lst` and `constr.lst` must be compatible, otherwise an error will be raised (see Karen documentation for details).

The following R code chunk shows how to fit a Kalman reaction network on the clonal tracking dataset previously simulated in Section S.7.1

```
nProc <- 1 # number of cores
cat(paste("\tLoading CPU cluster...\n", sep = ""))
cat(paste("Cluster type: ", "PSOCK\n", sep = ""))
cpu <- Sys.getenv("SLURM_CPUS_ON_NODE", nProc) ## define
    ↪ cluster CPUs
hosts <- rep("localhost",cpu)
cl <- makeCluster(hosts, type = "PSOCK") ## make the cluster
rm(nProc)

## mean vector of the initial condition:
m_0 <- replicate(nCL, X0, simplify = "array")
colnames(m_0) <- 1:nCL
## covariance matrix of the initial condition:
P_0 <- Diagonal(length(ctps) * nCL, 1e-5)
rownames(P_0) <- colnames(P_0) <- rep(1:nCL, each = length(
    ↪ ctps))
## Fit Karen on the simulated data:
res.fit <- get.fit(rct.lst = rcts,
                   constr.lst = cnstr,
                   latSts.lst = latsts,
                   ct.lst = ctps,
                   Y = XY$Y[,setdiff(ctps, latsts),],
                   m0 = m_0,
                   P0 = P_0,
                   cl = cl,
                   list(nLQR = 3,
                        lmm = 25,
                        pgtol = 0,
                        relErrfct = 1e-5,
                        tol = 1e-9,
                        maxit = 1000,
                        maxitEM = 10,
                        trace = 1,
                        FORCEP = FALSE))
stopCluster(cl) ## stop the cluster
```

##### S.7.3 Visualizing results

In this section we show how to visualize the main graphical output of Karen, that is the first two-order smoothing moments and the corresponding cell differentiation network. To this end we use the functions `get.sMoments()` and `get.cdn()` that receive as main input the result of a previously fitted Kalman reaction network provided by the function `get.fit()`. The function `get.sMoments()` receives also a 3-dimensional array for the stochastic process $X$ if available (see Karen documentation for details). The following R code shows how to obtain these graphics from the Kalman Reaction Network previously fitted in Section S.7.2.

```
## simulated data and smoothing moments
par(mar = c(2,5,2,2), mfrow = c(1,3))
get.sMoments(res.fit = res.fit, X = XY$X)
```

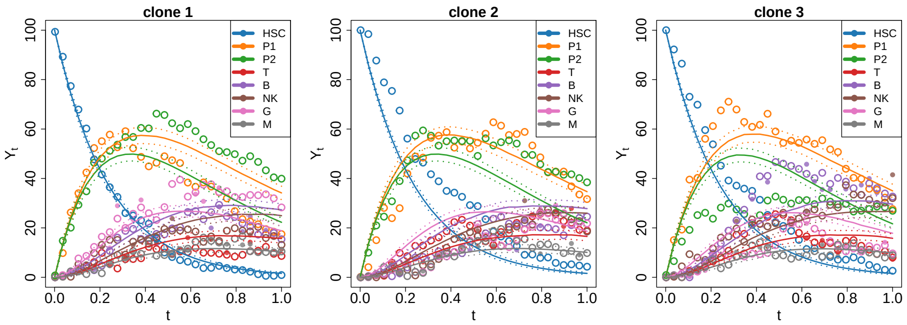


```
## Cell differentiation network
library(devtools)
source_url("https://raw.githubusercontent.com/jevansbio/
    ↪ igraphhack/master/igraphplot2.R")
environment(plot.igraph2) <- asNamespace('igraph')
environment(igraph.Arrows2) <- asNamespace('igraph')

legend_image <- as.raster(matrix(colorRampPalette(c("
    ↪ lightgray", "red", "black"))(99), ncol=1))
layout(mat = matrix(c(1,1,1,2), ncol = 1))
par(mar = c(0,0,3,0))
get.cdn(res.fit = res.fit,
        edges.lab = F)
plot(c(0,1),c(-1,1),type = 'n', axes = F,xlab = '', ylab = '
    ↪ ')
text(x=seq(0,1,l=5), y = -.2, labels = seq(0,1,l=5), cex =
    ↪ 2, font = 2)
rasterImage(t(legend_image), 0, 0, 1, 1)
```

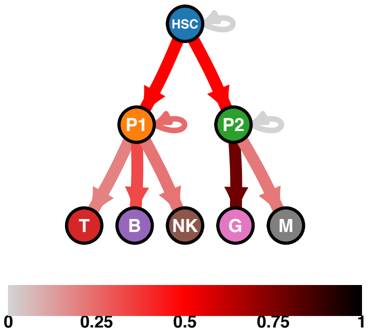


#### S.8 Data availability

The data that supports the findings of this study is openly available at [15–18].

#### S.9 Code availability

- The code that supports the findings of this study is openly available at `https://github.com/delcore-luca/CellDifferentiationNetworks`

- The stochastic framework is implemented in the R package Karen available at `https://github.com/delcore-luca/Karen`
